# PD-L1^+^ and Hyal2^+^ myeloid cells in renal cell carcinoma: a case report

**DOI:** 10.1101/2020.12.29.424690

**Authors:** Elizabeth Kwenda, Paul R. Dominguez-Gutierrez, Padraic O’Malley, Paul L. Crispen, Sergei Kusmartsev

## Abstract

RCC patients frequently have increased numbers of immunosuppressive myeloid cells in circulation. High numbers of myeloid derived suppressor cells (MDSCs) in the blood are associated with immune suppression as well as with cancer-related inflammation which drives the mobilization of myeloid cells to tumor tissue. Here we show that peripheral blood from a previously untreated renal cell carcinoma patient has increased numbers of monocytic CD33^+^CD11b^+^ MDSCs, which also co-expressed PD-L1 and membrane-bound enzyme hyaluronidase 2 (Hyal2). PD-L1 expression is associated with immune suppression, whereas expression of Hyal2 is associated with inflammation, because Hyal2^+^ myeloid cells can degrade the extracellular hyaluronan (HA), leading to the accumulation of pro-inflammatory HA fragments with low molecular weight. These findings implicate the potential involvement of monocytic MDSCs in both tumor-associated immune suppression and cancer-related inflammation. Analysis of organoid-like tumor-tissue slice cultures prepared from cancer tissue of the same patient revealed the significant presence of PD-L1^+^ HLA-DR^+^ macrophage-like or dendritic cell-like antigen-presenting cells in tumor stroma. Interestingly that stroma-associated PD-L1^+^ cells frequently have intracellular hyaluronan. Collectively, data presented in this study suggest that the interplay between tumor-recruited myeloid cells and stromal hyaluronan may contribute to the inflammation and immune tolerance in cancer.

## Introduction

Immune checkpoint inhibitors have improved the treatment of a broad spectrum of cancers including renal cell carcinoma (RCC), metastatic melanoma, and non-small lung cancer. These humanized monoclonal antibodies target inhibitory receptors (e.g. CTLA-4, PD-1, LAG-3, TIM-3) and ligands (PD-L1) expressed on T lymphocytes, antigen-presenting cells, and tumor cells and elicit an anti-tumor response by stimulating the immune system. However, both cancer-related inflammation and tumor-associated immune suppression frequently override the anti-tumor immune response. We previously reported that tumor-associated myeloid cells in RCC patients exert a tolerogenic function since they can up-regulate the expression of FOXP3 and CTLA-4 in autologous T lymphocytes (**1)**. These molecules are implicated in the mechanism of T cell unresponsiveness. Tumor associated macrophage (TAM)-mediated induction of FOXP3 and CTLA-4 expression in T lymphocytes favors tolerogenic conditions that allow tumors to escape the immune response. Also, the immunosuppressive ligand PD-L1 is frequently expressed in RCC and is associated with poor prognosis **(2**). Interestingly, PD-L1 expression can be detected in both myeloid cells and cancer cells (**3–6**). Here we show an increased presence of PD-L1^+^ myeloid cells in both peripheral blood and tumor-tissue from a previously untreated patient with clear cell RCC. Furthermore, tumor-infiltrating PD-L1^+^ myeloid cells express a marker of antigen-presenting cells HLA-DR and frequently show significant amounts of internalized hyaluronan, indicating the possible contribution of stroma and tumor-associated hyaluronan in the modulation of immune function of myeloid cells including antigen-presentation.

## Materials and Methods

### Human subjects

Freshly excised kidney tumor tissue and peripheral blood during radical nephrectomy from a previously untreated patient diagnosed with RCC. Clinical samples were collected from the patient after obtaining written informed consent. All samples were obtained according to federal guidelines and as approved by the University of Florida institutional review board (IRB).

### Reagents and culture medium

Biotinylated hyaluronan-binding protein (HABP) was supplied by Millipore-Sigma. Streptavidin conjugated with PE or FITC was purchased from Biolegend (San Diego, CA). Hyaluronidase-2 polyclonal antibody conjugated with PE or Alexa-488 was obtained from Bioss Antibodies. Anti-PD-L1 and other antibodies were acquired from Biolegend. *In vitro* experiments were conducted using complete culture media consisting of RPMI 1640 medium supplemented with 20 mM HEPES, 200 U/ml penicillin, 50 μg/ml streptomycin (all from Hyclone), and 10% FBS from ATCC (Manassas, VA).

### Preparation of tissue slices from human tumor kidney tissue

The precision-cut tissue slices, 2-4 mm in diameter and 200-300-micron thick, were produced using a Compresstome Vibratome VF-300-0Z. After cutting, tissue slices were placed into 24-well cell culture plates in complete RPMI-1640 medium supplemented with 10% FBS and antibiotics and cultured at 37° C in a humidified CO_2_ incubator.

### Isolation of CD11b myeloid cells from peripheral blood of cancer patients

PBMCs were isolated by gradient density centrifugation using Lymphoprep (Accu-Prep, 1.077g/ml, Oslo, Norway). CD11b myeloid cells were purified from PBMCs by positive selection using the anti-CD11b microbeads and columns (Miltenyi Biotec). Briefly, cells were incubated with beads conjugated with anti-mouse CD11b and positively selected on LS columns. The viability of all recovered cells was 95%, as determined by trypan blue exclusion.

### Visualization of tumor-produced HA

Tumor tissue slices were cultured for 7-14 days in 24-well cell culture plates in a humidified CO_2_ incubator at 37° C to allow for the production of HA. At the end of incubation, tissue-produced HA was found settled at the bottom of the culture plate wells. To monitor and visualize the accumulation of tissue-produced HA fragments on the plastic surface, the tissue slices and culture medium were removed at different time points. The empty wells were washed with warm PBS and fixed with 4% formaldehyde for 30 min. After fixation, plate wells were washed with PBS containing 2% FBS and incubated overnight with biotinylated HA-binding protein (3 μg/ml, Calbiochem-EMD Millipore) at 4° C (**7**). The next day, after washing the wells with PBS containing 2% FBS, streptavidin-conjugated with fluorochrome was added to the wells and incubated for 30 min at 4°C. Plates were then washed with PBS and the bottoms of the wells were visualized using EVOS (Invitrogen) or Lionheart (Biotek Instruments) immunofluorescent imaging microscopes.

### Immunofluorescent microscopy

Immunofluorescent staining and analysis were performed according to the previously described protocol (**6**).

## Results and Discussion

Cancer patients, including patients with RCC, frequently have increased numbers of immunosuppressive myeloid cells such as MDSCs (**8–10**). Thus, MDSCs isolated from the blood of patients, but not from healthy donors, were capable of suppressing antigen-specific T-cell responses *in vitro* through the secretion of reactive oxygen species and nitric oxide upon interaction with CTL. More recently (**11**), we showed that monocytic MDSCs in patients with bladder cancer, in contrast to healthy donors, frequently co-expressed the enzyme hyaluronidase 2 (Hyal2).

To examine whether myeloid cells in RCC patients also expressed Hyal2, we have isolated CD11b^+^ myeloid cells from the peripheral blood of a previously untreated patient with renal cell carcinoma. First, we looked at the presence of CD33^+^ monocytic and CD15^+^ granulocytic MDSCs. Data presented in **Fig 1A** demonstrate the high levels of CD33 and relatively low expression of CD15 among blood-derived myeloid cells. Similar to human bladder cancer, CD33^+^ in a patient with RCC, also frequently-co-expressed membrane-bound enzyme Hyal2 (**Fig 1B**, left image). Furthermore, the Hyal2^+^ myeloid cells also co-expressed the immunosuppressive ligand PD-L1 (**Fig 1 C, D**).

**Figure 1.**
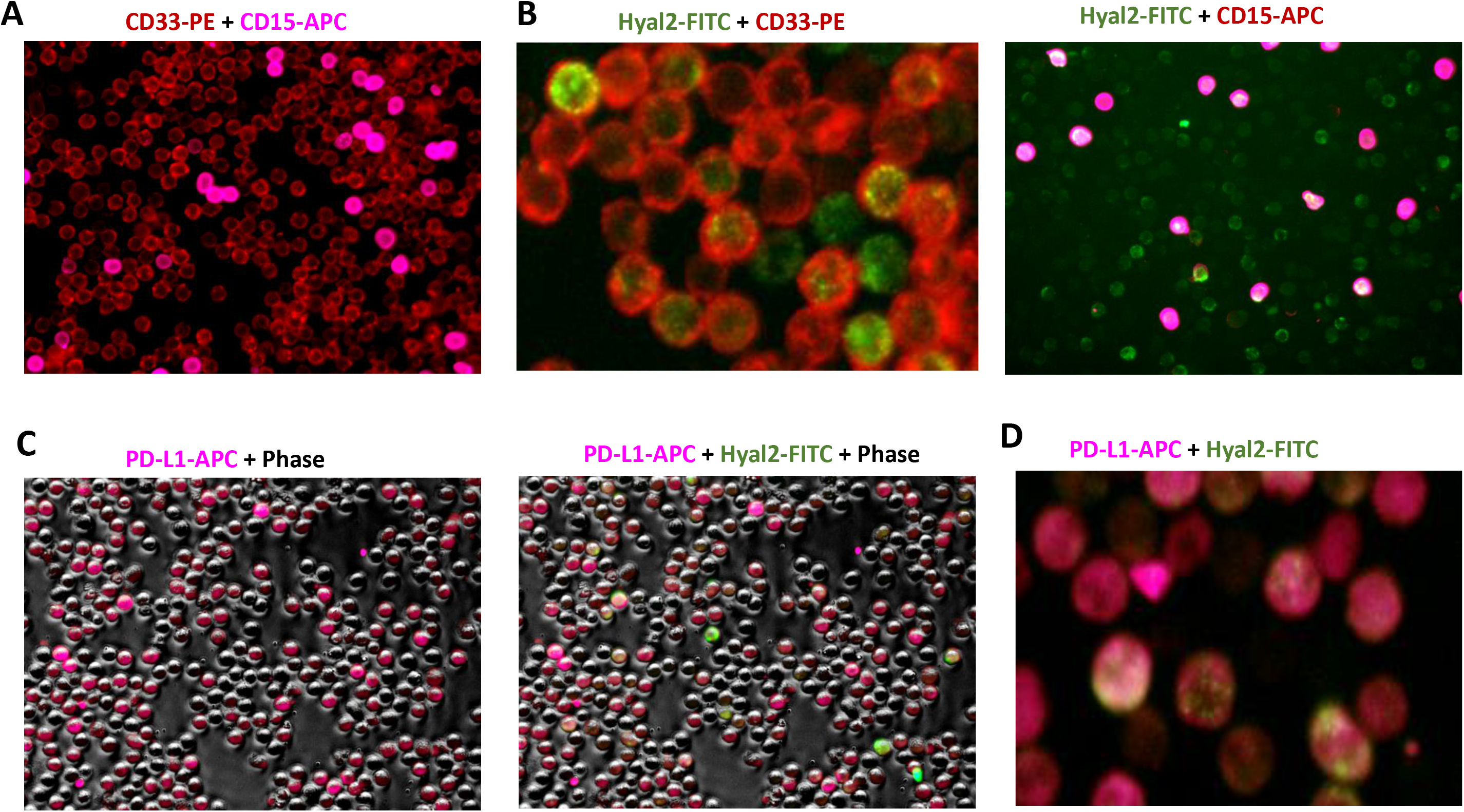
The increased presence of PD-L1 and Hyal2-expressing myeloid cell subsets in the patient’s peripheral blood. CD11b myeloid cells were isolated from the peripheral blood using magnetic beads. Freshly isolated cells were stained with CD33-PE, CD15-PE, and Hyal2-FITC antibodies (images **A, B**), or with PD-L1-APC and Hyal2-FITC antibodies (images **C, D**). Representative IF images are shown.

High numbers of MDSCs in the blood of cancer patients are frequently associated with cancer-related inflammation which drives the mobilization of myeloid cells to tumor tissue. To examine the tumor-associated myeloid cells in RCC tissue, we prepared the organoid-like cancer tissue slices using freshly excised tumor tissue from the same patient. Live imaging of stoma in RCC tissue slice cultures before fixation showed a significant presence of both irregularly shaped fibroblast-like large cells, and smaller round shaped macrophage-like or dendritic cell-like cells (**Fig. 2 A-D**). Moreover, the smaller cells frequently were observed in the close proximity of fibroblast-like cells **(Fig. 2 B)**, suggesting the potential interaction between those cells.

**Figure 2.**
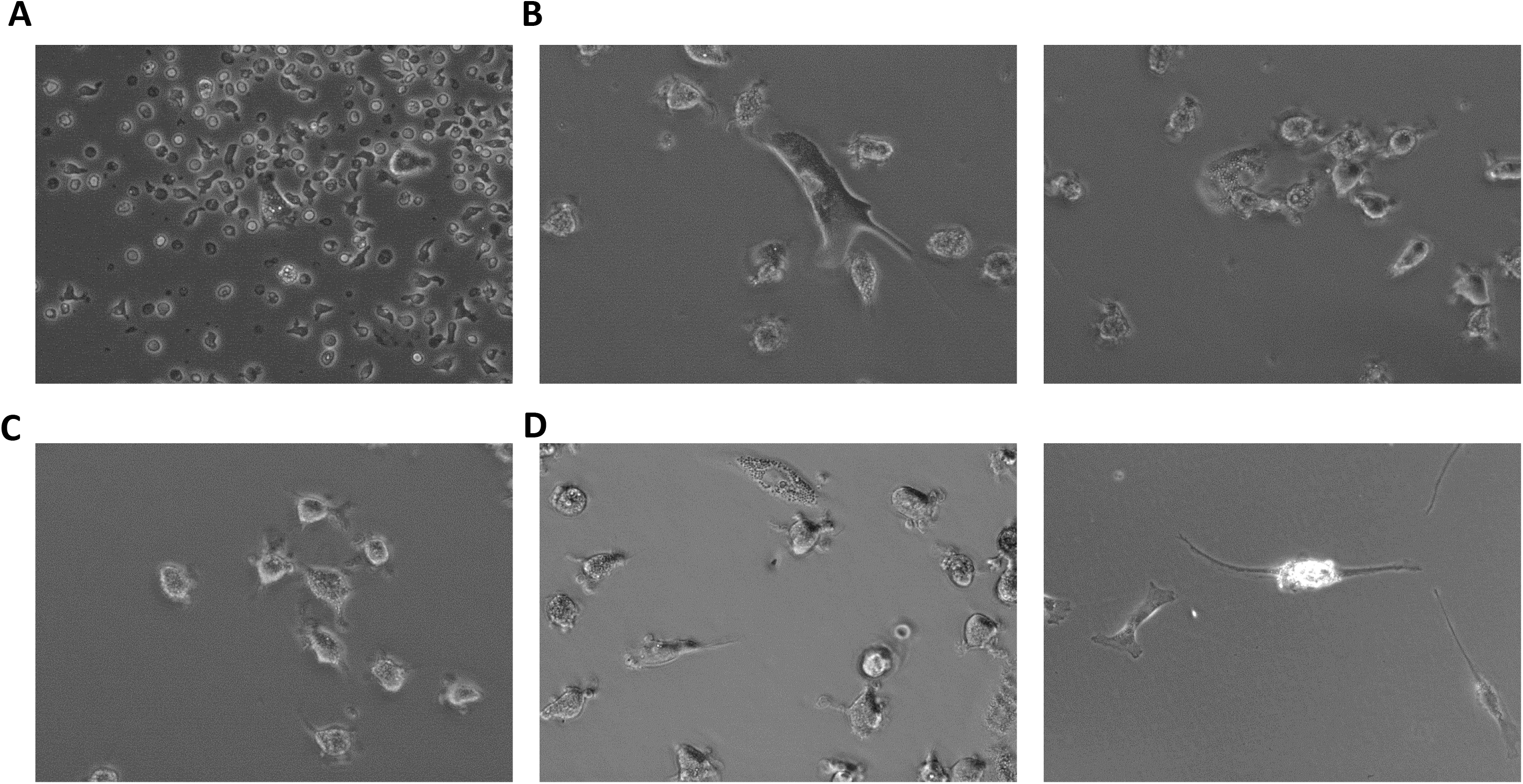
Tumor-infiltrating immune cells interact with cancer-associated fibroblast-like cells. Live imaging (before fixation). Representative bright-field images of tumor stroma from the same patient are shown (images **A-D**).

After fixation and washing of tumor tissue slices with PBS, we stained the remaining adherent stromal cells for the PD-L1. Data presented in **Fig. 3 A, B** demonstrate that majority of macrophage-like or dendritic-cell-like cells in RCC stroma express PD-L1. Co-expression of HLA-DR by these cells (**Fig. 3A**) supports the idea that these PD-L1^+^ cells belong to the antigen-presenting cells. Also, staining of RCC stroma for hyaluronan (HA) revealed the PD-L1^+^ cells have a marked presence of intracellular HA (**Fig. 3B-D**).

**Figure 3.**
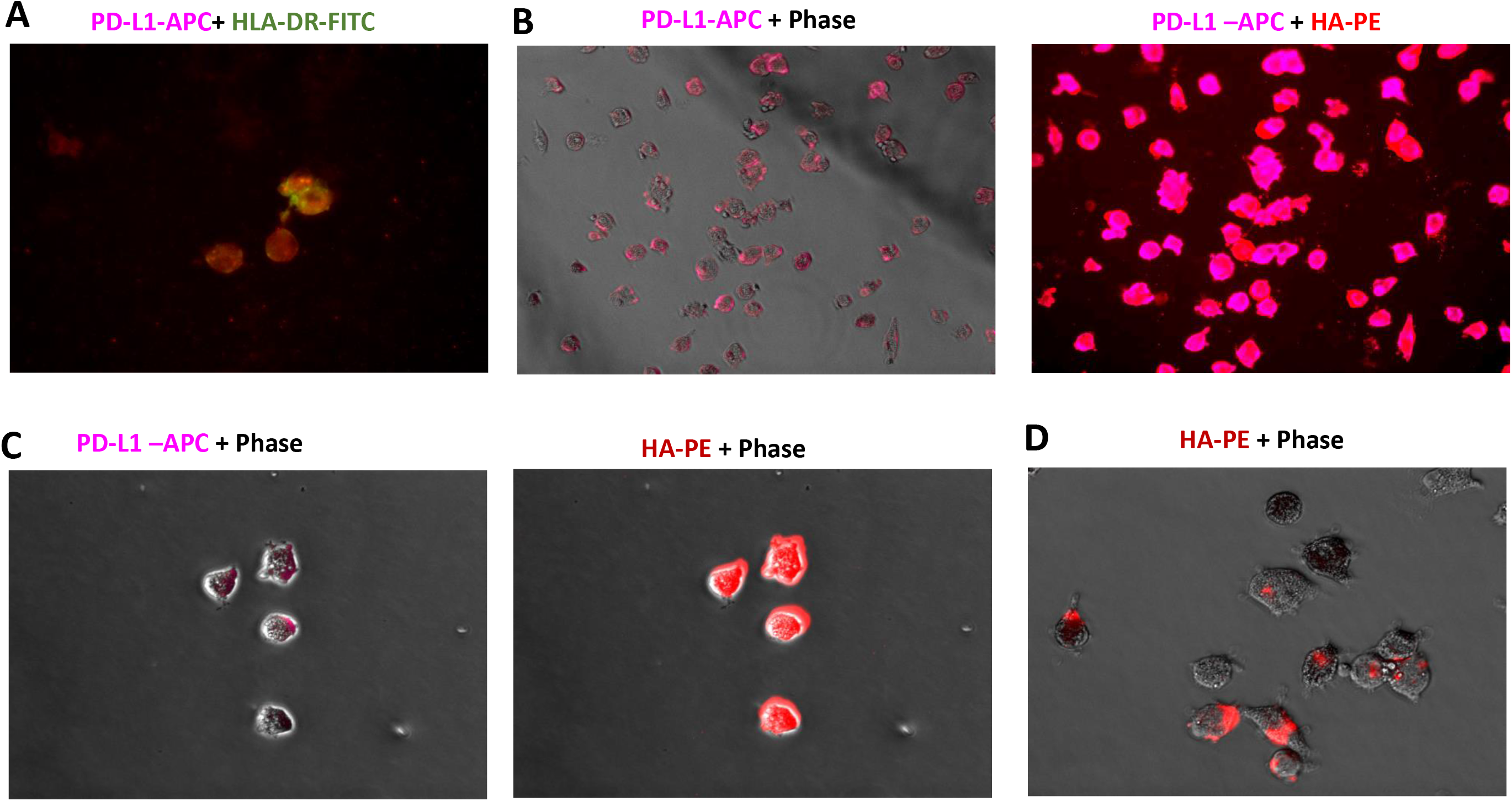
Visualization of intracellular HA in tumor-infiltrating PD-L1+ myeloid cells. The human cancer tissue slices were cultured for 7 days. Non-adherent cells were carefully removed from the plate. Plate with remaining adherent cells washed with PBS and fixed with 4% formaldehyde. Plate-bound cells were stained with HLA-DR-FITC (image **A**) and PD-L1-APC (images **A, B, C)** antibodies. To visualize the tumor-produced HA, biotinylated HA-binding protein and PE-labeled Streptavidin were subsequently added (images **B, C, D**). Representative images are shown.

Additional analysis revealed that stromal fibroblast-like cells expressed fibroblast-specific marker FAP-alpha (**Fig.4A**). These data are consistent with previous reports demonstrating the presence of FAP-alpha cancer-associated fibroblasts in RCC tissues (**12, 13**). Staining of stroma for the HA showed (**Fig. 4 B-C**) that localization of cancer-associated fibroblasts (CAFs) in RCC stroma is associated with HA (red) suggesting that CAFs contribute to the HA in the RCC tumor microenvironment.

**Figure 4.**
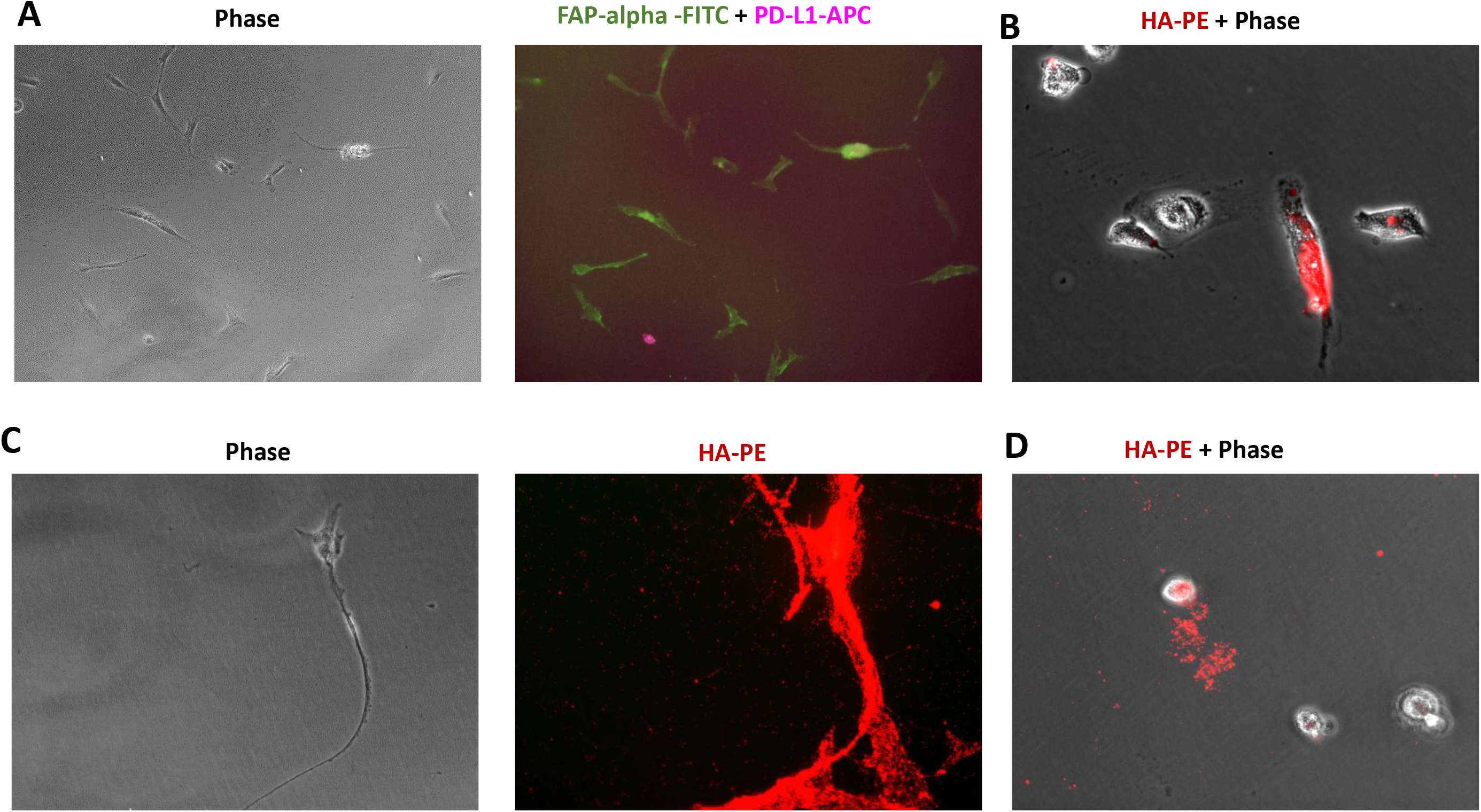
Detection of cancer-associated fibroblasts in kidney tumor stroma. The tissue slices were cultured for 7 days. Non-adherent cells were carefully removed from the plate. Plate with remaining adherent cells washed with PBS and fixed with 4% formaldehyde. Plate-bound cells were stained with FAP-alpha-FITC and PD-L1-APC (image **A)** antibodies. To visualize the tumor-produced HA, biotinylated HA-binding protein and PE-labeled Streptavidin were subsequently added (images **B, C, D**). Representative images are shown.

This is the second report confirming the increased mobilization of Hyal2+ myeloid cells in cancer patients following elevated numbers of Hyal2^+^ myeloid cells being detected in the peripheral blood of bladder cancer patients (**11**). Upon recruitment to the tumor, Hyal2^+^ expressing myeloid cells capable of degrading extracellular hyaluronan (HA) into small fragments, promoting the accumulation of HA fragments low molecular weight (20 kDa). Accumulation of LMW-HA fragments in tumor tissue has been associated with enhanced production of multiple inflammatory and pro-angiogenic factors. Since tumor stroma is rich in hyaluronan, it is plausible that tumor stroma involved in the regulation of an anti-tumor immune response. However, the exact mechanisms of stroma-immune interactions in cancers including RCC remain largely unknown. Our data demonstrate the frequent co-localization of HA and fibroblast-like cells in RCC stroma. The enhanced HA metabolism has previously been described in several major subtypes of RCC such as clear cell, papillary, and chromophobe renal carcinomas (**14**). Thus, the median transcript levels of hyaluronic acid synthase 1 (HAS1) and major HA receptors CD44 and RHAMM were elevated 3 to 25-fold in those tumor tissues when compared with normal tissues.

Here we show that peripheral blood of previously untreated RCC patient is enriched for the monocytic CD33^+^ MDSCs. These myeloid cells co-express immunosuppressive ligand PD-L1 as well as membrane-bound enzyme Hyal2. Expression of PD-L1 suggests involvement in the mechanisms of suppression (**15**), whereas Hyal2 expression indicates the involvement of these cells in process of HA degradation which contributes to cancer-related inflammation (**6, 16–18**). Upon recruitment to the tumor tissue, tumor-associated MDSCs frequently differentiate into immunosuppressive antigen-presenting cells such as macrophages (**19**). Our data indicate that macrophage-like HLA-DR^+^ cells in RCC stroma frequently express immunosuppressive PD-L1 ligand and also show significant amounts of internalized tumor-associated HA. Both tumor epithelial cells and CAFs were recently identified as the main sources of HA in the tumor microenvironment (**20**). Taken together, this work demonstrates that stromal HA metabolism in the tumor microenvironment likely contributes to the regulation of anti-tumor immune response through modulation of tumor-infiltrating APCs.

